# High-Intensity Interval Training Decreases Circulating HMGB1 in Individuals with Insulin Resistance; Plasma Lipidomics Identifies Associated Cardiometabolic Benefits

**DOI:** 10.1101/2024.08.21.608998

**Authors:** Gabriela Martinez Bravo, Prabu Paramasivam, Gabriella F. Bellissimo, Quiteria Jacquez, Huayu Zheng, Fabiano Amorim, Roberto Ivan Mota Alvidrez

**Affiliations:** Pharmaceutical Sciences, College of Pharmacy, University of New Mexico, Albuquerque, NM, US; Biomedical Engineering Department, University of New Mexico, Albuquerque, NM, US; Clinical and Translational Sciences Center, University of New Mexico, Albuquerque, NM, US; Health, Exercise and Sports Sciences, University of New Mexico, Albuquerque, NM, US; Cardiovascular and Metabolic Diseases (CVMD) Signature Program, University of New Mexico, Albuquerque, NM, US; Autophagy, Inflammation, Metabolism CoBRE, University of New Mexico, Albuquerque, NM, US

**Keywords:** HMGB1, Exercise, Insulin Resistance, HIIT, BW-HIIT, Inflammation

## Abstract

**Background:** Exercise is a fundamental primary standard of care for cardiometabolic health. Body Weight (BW) High-Intensity Interval Training (HIIT) is an effective strategy for reducing cardiometabolic markers in individuals with insulin resistance and Type-2 diabetes (T2D). High-mobility group box 1 (HMGB1), a ubiquitous nuclear factor, plays an ample role beyond an alarmin in T2D development and progression. Our group has described this novel role previously, showing the beneficial effect of whole body HMGB1 silencing in decreasing hyperglycemia in diabetic mice. In the present study we tested the hypothesis that BW-HIIT as an effective exercise training modality will decrease cardiometabolic risk with a concomitant decrease in circulating HMGB1 more prominently in insulin resistant individuals compared to non-insulin resistant individuals contrasting to what we can evidence in a preclinical murine model of insulin resistance; **Methods:** Human and mouse pre- and post-exercise serum/plasma samples were analyzed for Lipidomics as well as Metabolic and Cytokine Multiplex assays. Standard of care, as well as cardiometabolic parameters, was also performed in human subjects; **Results:** insulin resistant individuals had the most positive effect, primarily with a decrease in the Homeostatic Model Assessment of Insulin Resistance (HOMA-IR). as an index of insulin resistance as well as decreased HMGB1 post-exercise. Lipidomic analysis illustrated the highly beneficial effect of exercise training using a modified HIIT program, showing an enhanced panel of circulating lipids post-exercise exclusively in insulin resistant individuals. Plasma multiplex revealed significant translational heterogeneity in our studies with distinct metabolic hormone responses to exercise conditioning with a decrease in inflammatory markers in insulin resistant individuals; **Conclusions:** The current study demonstrated that 6-week BW-HIIT training improves cardiometabolic, anti-inflammatory markers, metabolic hormones, and insulin sensitivity in humans, strongly associated with decreased circulating HMGB1. Overall, these experiments reinforce the potential of HMGB1 as a marker of changes in insulin resistance and the positive effect of exercise training on insulin resistance possibly preventing the development of T2D and associated complications.

## 1. Introduction

Insulin resistance is a clinical condition where muscle, fat, and liver cells become less responsive to insulin, leading to impaired glucose uptake and elevated blood glucose levels [1]. The pancreas increases insulin production to compensate for high circulating glucose, but this compensatory mechanism eventually fails due to beta cell dysfunction [2]. Persistent hyperglycemia ensues, culminating in T2D when the pancreas can no longer produce sufficient and effective insulin to manage blood glucose levels effectively [3]. Contributing factors include genetic predisposition, obesity (visceral fat releases free fatty acids and inflammatory cytokines that disrupt insulin signaling), sedentary lifestyle, and diets based on highly refined carbohydrates and non-insulin resistant fats [4]. Early intervention through lifestyle changes such as improved diet and increased physical activity is crucial to prevent or delay the onset of T2D [3–5].

Exercise therapy is a cornerstone in the management of various chronic diseases, including T2D and inflammatory conditions known to benefit glucose control. However, it is fundamental to make standalone therapeutic targets that enhance the effect of exercise rather than exercise being just an adjuvant [6]. The therapeutic benefits of exercise are achieved through several physiological mechanisms that enhance overall health and reduce the risk of disease [7]. According to a systematic review by Shah et al. (2021), exercise interventions lasting over eight weeks led to significant reductions in glycated hemoglobin (HbA1c), fasting blood glucose levels, body mass index (BMI), and waist circumference. Exercise optimizes glycemic control and enhances the overall quality of life for individuals with T2D. Regular physical activity, including high-intensity exercises like HIIT and moderate exercises like walking, is recommended to complement medical treatments for effective diabetes management [8].

As mentioned, HIIT is regarded as an effective stimulus for preventing insulin resistance and reducing T2D risk [9]. HIIT involves short bursts of intense exercise followed by periods of rest or low-intensity exercise [10]. This training enhances insulin sensitivity possibly through several mechanisms: 1) HIIT increases glucose uptake by muscle cells by enhancing the activity of glucose transporter Type-4 (GLUT4), which facilitates glucose entry into the cells. 2) It improves mitochondrial function and increases the number of mitochondria in muscle cells, which enhances their capacity to oxidize glucose and fatty acids for energy. 3) HIIT promotes fat loss, particularly visceral fat, and is strongly associated with insulin resistance. 4) HIIT reduces inflammation and oxidative stress, which can impair insulin signaling pathways. These physiological changes collectively improve insulin sensitivity and blood glucose control [10, 11]. The systematic review by Shah et al. (2021) found that exercise interventions reduced HbA1c by 0.5% to 0.8% on average, indicating significant improvements in long-term glycemic control [8]. Similarly, Colberg et al. (2010) found that regular physical activity lowered HbA1c by approximately 0.7%, confirming the beneficial effect of exercise on blood sugar levels [7]. Another example of exercise improvement is demonstrated in the paper by Balducci et al. (2010), which shows how aerobic exercise improved HOMA-IR by 20-30%, indicating enhanced insulin sensitivity [12]. Way et al. (2016) Reported similar states showing an improvement of 25% in HOMA-IR following a combination of aerobic and resistance training [13].

A key factor in the regulation and progression of insulin resistance is the nuclear protein High-mobility group box 1 (HMGB1). This active protein is particularly significant during oxidative stress and inflammation as it triggers various inflammatory signaling pathways, including the toll-like receptors (TLRs) and receptors for advanced glycation end products (RAGE) [14]. The activation of these receptors leads to the production of pro-inflammatory cytokines, such as TNF-α and IL-6, which disrupt insulin signaling pathways [14]. Another contributing factor is the Reactive Oxygen Species (ROS) generated during oxidative stress, which damages cellular components and hampers insulin receptor function, thereby worsening insulin resistance [14, 15]. Another role of HMGB1 is activating immune cells and regulating insulin secretion by influencing pancreatic β-cell function. Excessive HMGB1 levels can impair insulin secretion, whereas controlled levels are necessary for optimal β-cell function and insulin release, but a person with T2D has higher levels of HMGB1[16]. HMGB1 acts as a pro-inflammatory cytokine that contributes to the development of many diabetic complications [17]. We hypothesize that regular physical activity reduces HMGB1 levels, alleviating its pro-inflammatory effects and improving insulin sensitivity in the cells. This contributes to restoring glucose homeostasis and better glycemic control in diabetic individuals. HMGB1 has become a significant biomarker in understanding glucose metabolism and the inflammatory processes associated with T2D and other medical conditions due to its involvement in inflammation and tissue damage [18]. Traditional biomarkers for T2D focus on blood glucose levels, HbA1c, and insulin sensitivity [19]. However, these often overlook chronic inflammation, critical in insulin resistance and glucose dysregulation. HMGB1 is unique in its potential as a biomarker for diagnosing inflammatory conditions, assessing disease severity, and predicting patient outcomes [18]. Additionally, the acetylated form of HMGB1 (acetyl-HMGB1) is a modified form actively secreted by cells in response to stress and inflammation. This form is particularly potent in signaling and modulating immune responses, further influencing glucose metabolism. It has gained attention for its distinct functions and potential as a biomarker, further expanding medicine’s diagnostic and prognostic capabilities [20]. In conclusion, HMGB1 and its variants are promising for improving disease management and advancing personalized medicine, emphasizing the importance of glucose control and inflammation in biomarker research.

Our group’s research has yielded highly promising results in recent years in the role of HMGB1 in insulin resistance and T2D. We have advanced our knowledge of HMGB1 beyond an alarmin. We have discovered that inhibiting HMGB1 activity in mice not only improves insulin sensitivity and reduces hyperglycemia but also underscores the potential of HMGB1 as a therapeutic target in T2D disease progression. These findings will pave the way for future treatments for insulin resistance. In this study, we delve into the effects of BW-HIIT and its role in reducing circulating HMGB1, offering a potential solution to this health issue [21].

## 2. Materials and Methods

### 2.1 Human exercise model

For this study, all individuals were informed of their enrolment’s objectives, procedures, potential risks, discomforts, and benefits. Informed consent was obtained before participation. The University of New Mexico (UNM) Institutional Review Board (IRB) Main Campus approved this study with reference number 21-319 to Dr. Fabiano.

Fourteen individuals (2 male/12 female) enrolled in the study meeting the following inclusion criteria: 11) adults ages 18 to 55 years, 2 2) classified with obesity (BMI ≥ 30 kg/m, ²), but otherwise non-insulin resistant, 3) physically inactive, individuals meeting less than 150 minutes of moderate to vigorous intensity physical activity per week, and 4) nonsmokers. The exclusion criteria were individuals who reported taking glucose or lipid lowering medications; unknown/potential prediabetes or T2D determined by aa fasting ((≥8hrs) blood glucose sample, analyzed by a portable glucose meter (TRUE METRIX Blood Glucose MonitorTM, Trividia Health, Inc., USA. Before starting the study, blood glucose levels were checked by sending samples to a commercial lab (Quest Direct™, Albuquerque, NM, USA) for fasting blood glucose and hemoglobin A1c (HbA1c).

Baseline and post testing procedures were performed at the UNM Exercise Physiology laboratory. Individuals were instructed to arrive in a hydrated and fasted state for ≥8 hours and to abstain from alcohol and vigorous exercise for 24 hours. Before body composition measurement, hydration status was checked using urine specific gravity (USG), and individuals with >1.020 were considered dehydrated. Next, individuals were provided a Fitbit Inspire 3 wearable activity tracking (WAT) device, which is an electronic monitoring devices that enable users to track health-related fitness metrics such as steps, activity level, walking distance, heart rate, and sleep patterns [22]. And instructed to install the Fitbit application on their smartphones. After the WAT was set up, individuals completed maximal voluntary isometric strength tests and treadmill peak oxygen consumption (_2_peakVO_2_peak) tests. Post-testing data collection procedures were the same as the baseline testing apart from the glucometer check and setting up the individual’s WAT device. All excluded individuals were provided with their results and advised to seek further medical screening. Fasting blood samples of approximately 15 mL were collected from the antecubital vein. Samples were collected in anticoagulant treated EDTA tubes for plasma separation or in untreated tubes to allow the blood to clot by leaving it undisturbed at room temperature. Serum was obtained by centrifuging the tubes for 15 min (1000g, 22°C) (Allegra X-14R Centrifuge, Beckman Coulter, Brea, CA) and stored at -80°C for subsequent analysis. Pre- and post-intervention quantitative measurements of leptin (Crystal Chem High Performance Assays, Inc., Illinois, USA) and IL-6 (Quantikine R&D System, Minneapolis, USA) were assessed using enzyme-linked immunosorbent assays (ELISA) kits. ELISA kit procedures were performed according to the manufacturer’s instructions, and the average intraassay coefficient of variations was 2.9% for leptin and 24% for IL-6. Serum concentrations of insulin, glucose, low-density lipoprotein (LDL), high-density lipoprotein (HDL), triglycerides (TG), total cholesterol (TC), and hs-CRP, and the EDTA sample for hemoglobin A1c (HgbA1c) were sent to a commercial laboratory (Quest Direct™, Albuquerque, NM, USA). Insulin resistance was calculated using HOMA-IR (fasting insulin (uIU/mL) x fasting glucose (mg/dL)/405) [23, 24].

After all baseline procedures were performed, individuals were instructed how to perform the remote exercise program. The program involved maximal effort Body Weight High Intensity Interval Training (BW-HIIT), prescribed for three days a week over 6 weeks. The BW-HIIT workouts were YouTube-based, consisting of a warm-up, two sets of five intervals, and a two-minute rest period after the first five intervals. The progression of the videos was every two weeks involving an increase in the duration of the maximal effort work interval and a decrease in the duration of the recovery interval. The YouTube BW-HIIT-based videos are available at: https://youtube.com/playlist?list=PLpqCdMYQbSNmA8zmjzuo2eVFj-IpQmFS_.. During the 6 weeks of remote training, individuals were provided an exercise journal to record their overall rating of perceived exertion (RPE) immediately after each workout. Additionally, participants individuals recorded their heart rate after each maximal effort interval using their WAT device. The WAT was also used to assess compliance with the program by evaluating physical activity (PA) minutes on users’ Fitbit profiles [25]. After the 6-week period, the individuals returned to the lab, where we performed the same analysis as the baseline pre-exercise visit.

### 2.2 Animal model of exercise training

All animal experiments were approved by the ethics committee of the University of New Mexico (UNM) Institutional Animal Care and Use Committee (IACUC) and Animal Welfare Committee. All live animal studies were conducted ethically, following relevant guidelines and regulations at the University of New Mexico under protocol number 23-201405-HSC. All animals were housed under the Animal Resource Facilities (ARF) at UNM. Adult male mice (5 per group) were used at 10 weeks of age under 2 groups: sedentary controls, meaning that they had normal diets and no exercise, and the other group, the exercised mice, meaning that they ran for 1hr on treadmills at 80 meters/min for 4 weeks. They were all on a normal diet. All exercise mice were acclimated for 1 week before starting the study, meaning they were placed in the treadmills for more time until they were comfortable running at the set speed. We collected blood at baseline, meaning at 10 weeks of age, and then at the end of the age-matched exercise period (14 weeks of age for sedentary mice). At the end of the experimental protocol, the animals were euthanized using a dose of 0.01 mL/g of Ketamine/Xylazine. Tissues were harvested including whole blood for serum and plasma isolation, aorta, liver, and muscle were harvested.

### 2.3 Human and murine cytokine and metabolic blood panel analysis

Human pre- and post-exercise serum were sent to Eve Technologies Corporation to perform the Human Metabolic Hormone 12-Plex Discovery Assay. This assay uses Luminex xMAP technology for multiplexed quantification of 12 Human cytokines, chemokines, and growth factors. The multiplexing analysis was performed using the Luminex™ 200 system (Luminex, Austin, TX, USA) by Eve Technologies Corp. (Calgary, Alberta). According to the manufacturer’s protocol, twelve markers were simultaneously measured in the samples using Eve Technologies’ Human Metabolic Hormone 12-Plex Discovery Assay® (MilliporeSigma, Burlington, Massachusetts, USA). The 12-plex consisted of Amylin(active), C-Peptide, Ghrelin, GIP, GLP-1(active), Glucagon, Insulin, Leptin, MCP-1, PP, PYY, and Secretin. Assay sensitivities of these markers range from 0.6 – 46.9 pg/mL for the 12-plex. Individual analyte sensitivity values are available in the MilliporeSigma MILLIPLEX® MAP protocol.

And Human Cytokine Panel 4 12-Plex Discovery Assay, this study used Luminex xMAP technology for multiplexed quantification of 12 human cytokines, chemokines, and growth factors. The multiplexing analysis was performed using the Luminex™ 200 system (Luminex, Austin, TX, USA) by Eve Technologies Corp. (Calgary, Alberta). According to the manufacturer’s protocol, twelve markers were simultaneously measured in the samples using Eve Technologies’ Human Cytokine Panel 4 12-Plex Discovery Assay® (MilliporeSigma, Burlington, Massachusetts, USA). The 12-plex consisted of BAFF, BRAK, CXCL16, HCC-4, HMGB1*, IFNβ, IL-24, IL-28B, IL-35, IL-37, MIP-4, and YKL40. Assay sensitivities of these markers range from 2.0 - 3800 pg/mL for the 12-plex. Individual analyte sensitivity values are available in the MilliporeSigma MILLIPLEX® MAP protocol.

For mice, serum from the 2 groups was isolated and sent to Eve Technologies Corporation to perform the Mouse Metabolic Hormone 12-Plex Discovery Assay, this study used Luminex xMAP technology for multiplexed quantification of 12 Mouse cytokines, chemokines, and growth factors. The multiplexing analysis was performed using the Luminex™ 200 system (Luminex, Austin, TX, USA) by Eve Technologies Corp. (Calgary, Alberta). According to the manufacturer’s protocol, twelve markers were simultaneously measured in the samples using Eve Technologies’ Mouse Metabolic Hormone 12-Plex Discovery Assay® (MilliporeSigma, Burlington, Massachusetts, USA). The 12-plex consisted of Amylin(active), C-Peptide 2, Ghrelin, GIP (total), GLP-1(active), Glucagon, Insulin, Leptin, PP, PYY, Resistin and Secretin. Assay sensitivities of these markers range from 1.4 – 91.8 pg/mL for the 12-plex. Individual analyte sensitivity values are available in the MilliporeSigma MILLIPLEX® MAP protocol.

Mouse Cytokine 32-Plex Discovery Assay was performed using Luminex xMAP technology for multiplexed quantifying 32 Mouse cytokines, chemokines, and growth factors. The multiplexing analysis was performed using the Luminex™ 200 system (Luminex, Austin, TX, USA) by Eve Technologies Corp. (Calgary, Alberta). According to the manufacturer’s protocol, thirty-two markers were simultaneously measured in the samples using Eve Technologies’ Mouse Cytokine 32-Plex Discovery Assay® (MilliporeSigma, Burlington, Massachusetts, USA). The 32-plex consisted of Eotaxin, G-CSF, GM-CSF, IFNγ, IL-1α, IL-1β, IL-2, IL-3, IL-4, IL-5, IL-6, IL-7, IL-9, IL-10, IL-12(p40), IL-12(p70), IL-13, IL-15, IL-17, IP-10, KC, LIF, LIX, MCP-1, M-CSF, MIG, MIP-1α, MIP-1β, MIP-2, RANTES, TNFα, and VEGF. Assay sensitivities of these markers range from 0.3 – 30.6 pg/mL for the 32-plex. Individual analyte sensitivity values are available in the MilliporeSigma MILLIPLEX® MAP protocol.

### 2.4 Metabolomics analysis

Targeted metabolomics analysis of plasma samples was conducted using MxP® Quant 500 kit (Biocrates, Innsbruck, Austria), in which 630 metabolites from 26 biochemical classes were assessed. Sample preparation, data acquisition method, and data processing followed the manufacturer’s protocols. In brief, 10μL of blank solution (Phosphate buffered saline), calibration standard solutions, quality control solutions, and human plasma samples were added into a 96-well plate, which was pre-incorporated with internal standards provided by Biocrates. The plate was dried under nitrogen flow using a positive pressure manifold (Positive Pressure-96 Processor, Waters Corporation, Milford, MA, USA). Then, a derivatization solution containing 5 % phenyl isothiocyanate was added to all the wells, and the plate was incubated for 60 min at room temperature (RT) followed by another drying step with the manifold. An extract solvent of 5 mM ammonium acetate in methanol was added to each well, followed by shaking at RT for 30 min. The extracts were eluted into a new 96-well plate using the manifold and further diluted with either ultrapure water or flow injection analysis (FIA) solvent (provided with the Biocrates kit) for liquid chromatography-tandem mass spectrometry (LC-MS/MS) and FIA tandem mass spectrometry (FIA-MS/MS), respectively. The LC-MS/MS and FIA-MS/MS measurements were performed to quantify small molecules and lipids, respectively, using an ExionLCTM UHPLC system coupled to a 5500 QTRAP® a triple quadrupole mass spectrometer (AB Sciex, Darmstadt, Germany). The LC-MS/MS at positive and negative ion modes was run first. The LC column was an MxP® Quant 500 kit system column system (provided with the Biocrates kit). Mobile phase compositions, temperature, and LC gradient elution conditions were set following the methods from Biocrates. FIA-MS/MS test was run after removing the column. The data were acquired using the Analyst (Sciex) software with multiple reaction monitoring (MRM) for all metabolites with optimized MS parameters provided by the manufacturer. WebIDQ cloud software (Biocrates, Innsbruck, Austria) was used for data processing to validate, quantify, and export data. The concentrations (μM) for each metabolite in different classes were calculated using WebIDQ. MetaboINDICATOR provided the relevant pathways and biological functions linked with metabolite sums and ratios as a built-in tool from WebIDQ.

### 2.5 Statistical analysis

Insulin resistance was calculated using HOMA-IR (fasting insulin (uIU/mL) x fasting glucose (mg/dL)/405) [26] and insulin resistant sub-analysis was performed regardless of sex but based on HOMA-IR. A sub analysis was evaluated then in pre- and post-exercise measurements. For cardiometabolic and metabolomics analysis, statistical analyses were performed using IBM SPSS (IBM Corp., Version 23.0 Armonk, NY, USA), except for Cohen’s dz, which was calculated using G*Power (G*power, Dusseldorf, Germany). All illustrated data was performed using GraphPad Prism v8.4 (GraphPad Software, Inc., USA). Unpaired two-tailed Student’s t-tests and one-way ANOVA were performed when appropriate and detailed in each section and each figure legend. Figure legends specify the number of replicates and samples analyzed. P-values <0.05 were determined to be significant. Data are presented as mean ± standard error (SE) unless otherwise indicated. The specific number of subjects is described for each assay. Statistical significance was set at *: <0.05, **: 0.001–0.01, ***: 0.0001–0.001, and ****: <0.0001.

## 3. Results

### 3.1 Circulating cytokine and chemokines panels post-exercise evidence heterogeneic effects in analysis between humans and mice exercise training

Plasma multiplex analysis demonstrated the longitudinal changes in circulating levels of cytokines, chemokines, and growth factors in mice and humans pre- and post-exercise (Fig. 2). Anti-inflammatory cytokines Il-4, IL-10 (p: 0.0471), IL-12p40 (p: 0.0443), IL-12p70 (p: 0.0009), IL-13, and MIP-1a were shown to be increased in exercise-conditioned mice. Pro-inflammatory cytokines IFNγ, IL-1α, IL-2, IL-5, IL-7, LIF, and LIX were evidenced to be reduced. Contrastingly, ILβ, IL-3 (p: 0.0542), IL-9, IL-15, IL-17, IP-10, KC, CSF, MIG, MIP-1b, MIP-2 (p: 0.0039), RANTES, TNFα, VEGF and the chemokine Eotaxin (p: 0.0043) were increased post-exercise in mice (Fig. 2A) (Unmatched graphs are shown in supplementary Fig. 1). In human samples HMBG1, BAFF, INFβ, BRAK, IL-28b, and YKL40 are increased post-exercise (Fig. 2B). HCC, IL-37, and CXCL16 levels decreased post-exercise (Fig. 2B). Importantly, results evidence heterogeneity in the pro-inflammatory cytokine, showing how IL-6 decreases in humans and mice post-exercise (Fig. 2C), and the chemokine MCP-1 (Fig. 2D).

**Fig. 1.**
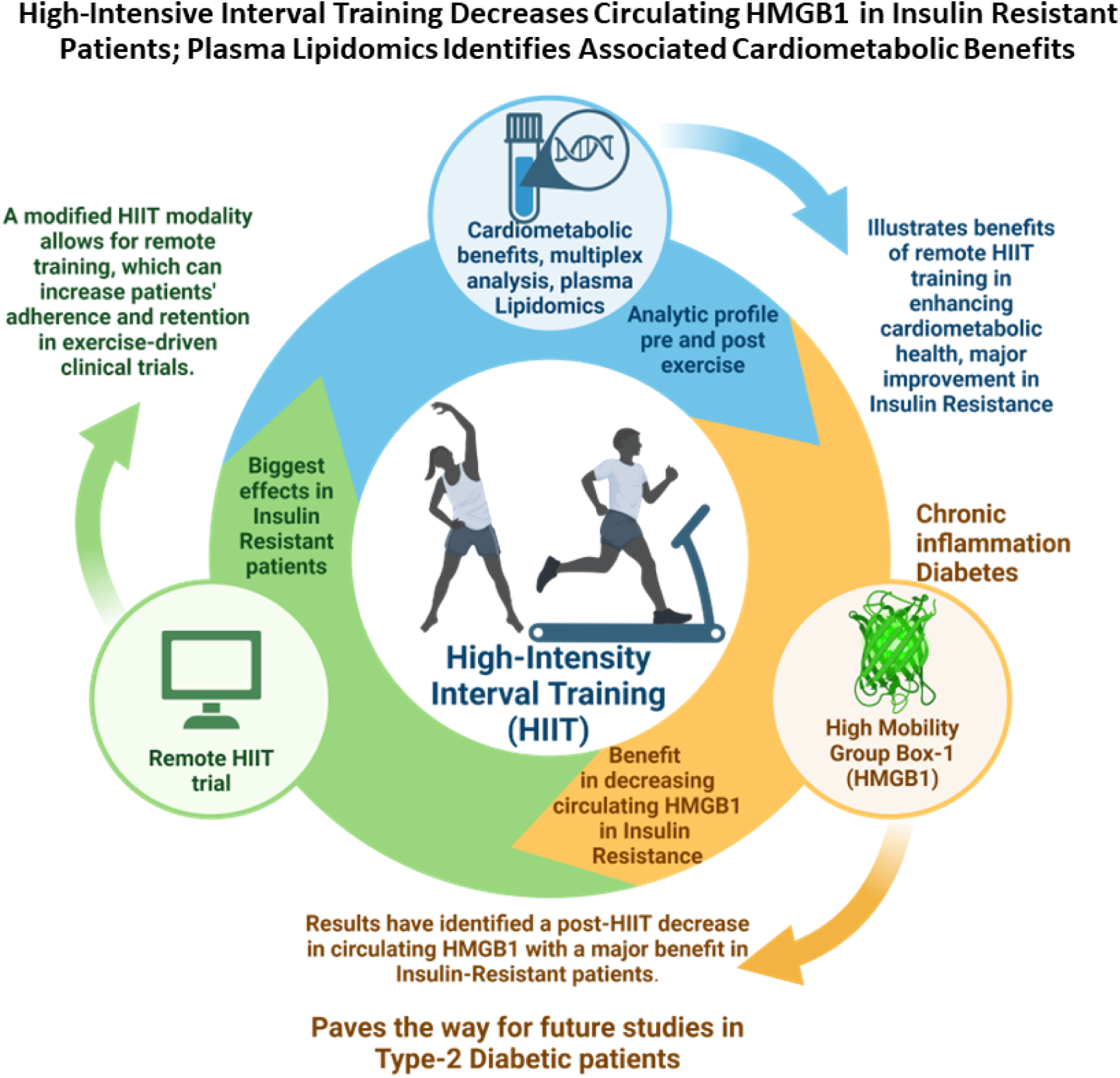
Graphical abstract. Created with BioRender

**Fig. 2.**
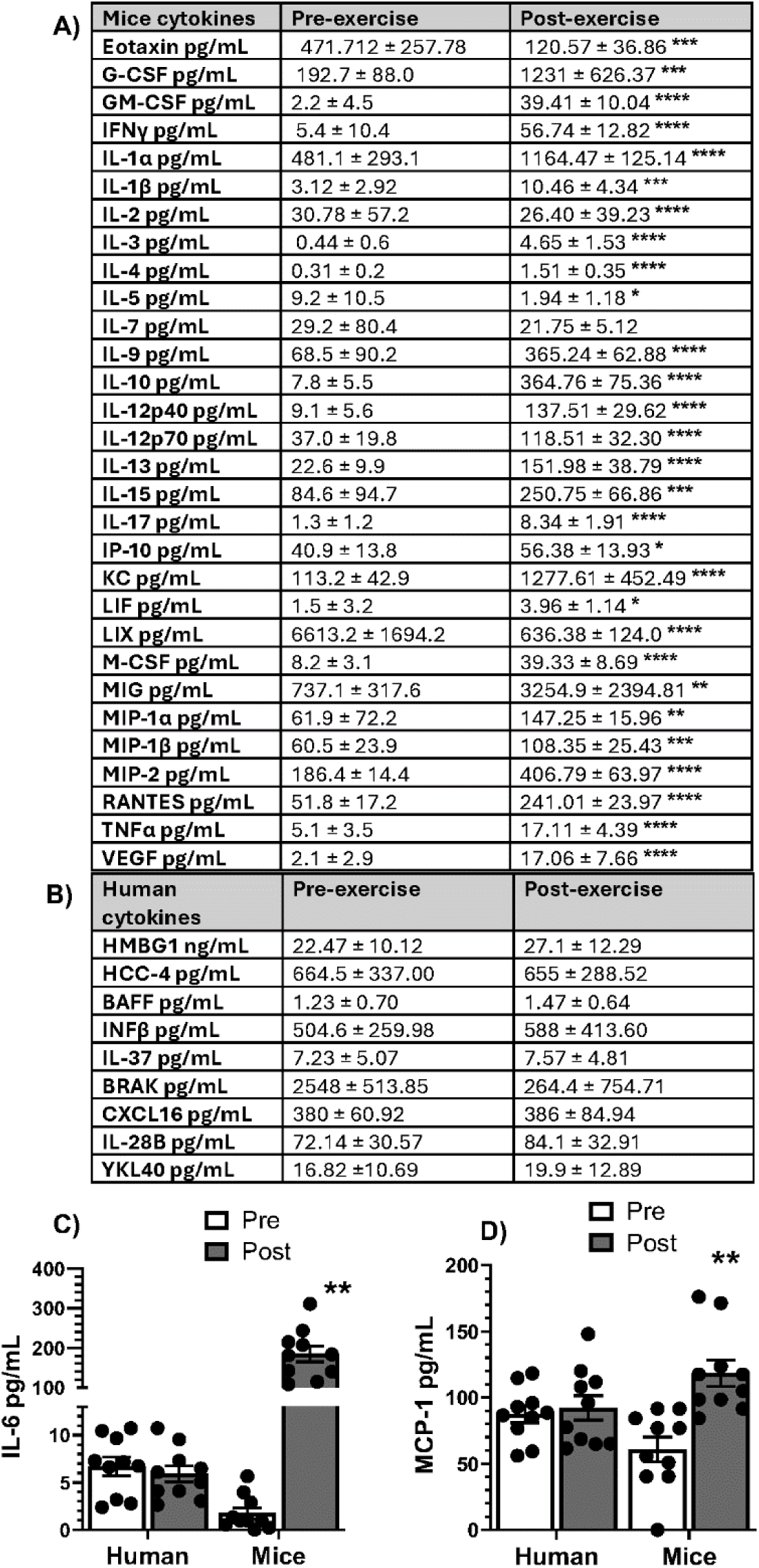
Plasma multiplex quantification analysis evidence exercise conditioning increases cytokines, chemokines, and growth factors, primarily in mice. Change of cytokine levels in humans after 16 weeks of exercise and 8 weeks of treadmill training in mice. A) Unmatched cytokines for mice and B) humans; C) IL6 levels and D) MCP-1 levels were assessed and matched in both humans and mice. All data represented mean+/-SEM. All cytokine levels in post-exercise compared to pre-exercise of humans (n=10) and mice (n=5) were used for independent student t-test analysis. Significance levels: *:<0.05, **: 0.001-0.01, ***: 0.0001, ****: <0.0001.

### 3.2 Metabolic and hormonal panels evidence distinct responses to exercise in humans and mice

Plasma multiplex was performed in humans and mice; the analysis revealed significant heterogeneity in the groups’ metabolic hormone responses to exercise conditioning. Both groups showed an increase in several hormones, indicating the positive impact of exercise in the gut, adipokine, and insulin hormone levels in BW-HIIT and exercise animals. Our findings varied between groups (Fig. 3); the hormones with an increase in both groups post-exercise are: Amylin with an increased average from 35.64 pg/mL to 50.0 pg/mL in humans, in mice the average increased from 53.76 pg/mL to 59.66 pg/mL (Fig. 3A); C-peptide with the increased average from 2,685.77 pg/mL to 2,948.26 pg/mL in humans, while in mice, the average increased from 157.61 pg/mL to 227.99 pg/mL (Fig. 3B); Glucagon with humans showing an increase average from 36.65 pg/mL to 41.53 pg/mL and mice from 22.49 pg/mL to 45.81 pg/mL (Fig. 3C). and Glucagon Inhibitory Peptide 1 with humans showing an increase from 24.63 pg/mL to 30.915 pg/mL and mice from 30.79 pg/mL to 64.92 pg/mL (Fig. 3H). While hormones showing more heterogeneity are Pancreatic Peptide (Fig. 3D); Ghrelin (Fig. 3E); insulin, post-exercise levels were significantly higher in humans but showed no significant change in mice (Fig. 3F); Peptide YY (Fig. 3G); and Leptin showing a significant increase in mice, but no significant change in humans (Fig. 3I). Unmatched comparisons of 2 metabolic hormones in mice were presented, Secretin levels significantly increased post-exercise from 13.00 pg/mL to 23.82 pg/mL (Supplementary Fig. 2A), and Resistin (Supplementary Fig. 2B).

**Fig. 3.**
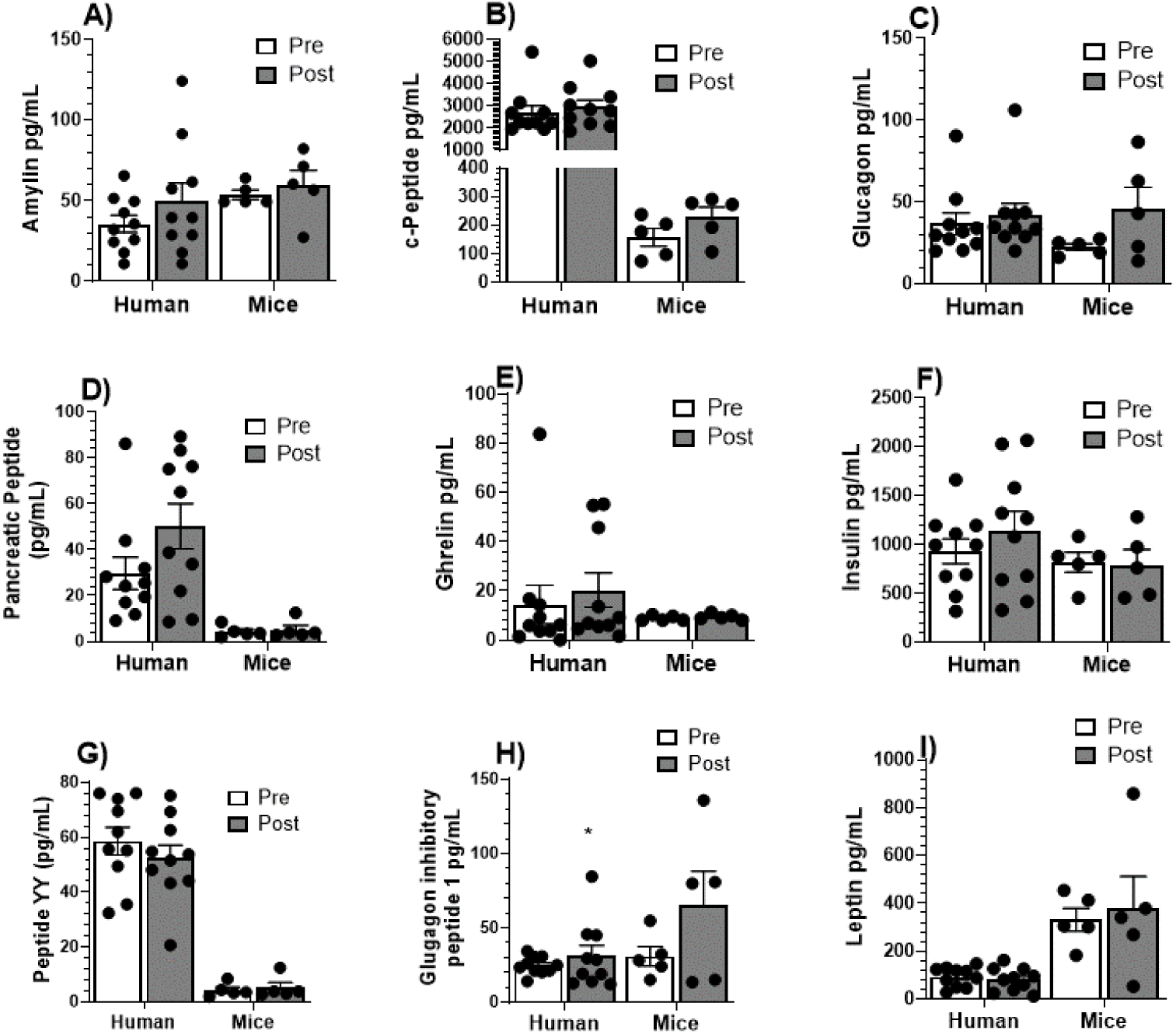
Plasma multiplex analysis evidence heterogeneity in metabolic benefits post-exercise between human and mouse exercise conditioning. Changes in metabolic hormone levels in humans after 16 weeks of exercise and 8 weeks of treadmill exercise training in mice. Matched comparisons are shown pre- and post-exercise in both humans and mice for: A) Amylin, B) c-peptide, C) Glucagon, D) Pancreatic peptide, E) Ghrelin, F) Insulin, G) Peptide YY, H) Glucagon inhibitory peptide 1 and I) Leptin. All data represented mean+/-SEM. All metabolic hormone levels in post-exercise compared to pre-exercise of humans (n=10) and mice(n=5) were used for independent student t-test analysis. Significance levels: *:<0.05, **: 0.001-0.01, ***: 0.0001, ****: <0.0001.

### 3.3 Reduced cardiometabolic risk factor in insulin resistant individuals involved in post-exercise

We evaluated the changes in cardiometabolic markers evidenced by the percentage of change from baseline (pre-exercise to post-exercise) in inflammatory markers (CRP and IL6), glycemic markers (HbA1c), and Leptin (marker useful in feeling full and satiety) to investigate the effects of exercise when we split our analysis to changes in non-insulin resistant vs. insulin resistant individuals. We determined insulin resistance using individuals with HOMA-IR < 2 were considered non-insulin resistant, while those with HOMA-IR > 2 were considered insulin resistant (Fig. 4). The results show reduced cardiometabolic risk factors post-exercise in individuals with insulin resistance compared to non-insulin resistant individuals. Cardiometabolic risk factors such as systolic blood pressure decreased -4.2% (Fig. 4D); cholesterol -2.8% (Fig. 4F); non-HDL -1.4 % (Fig. 4I), and cholesterol/HDL ratio -0.7 % (Fig. 4H) in insulin resistant individuals compared to non-insulin resistant individuals. At the same time, pro-inflammatory markers like C-reactive protein decreased by 6.60% (Fig. 4A) and IL6 by 0.02 % (Fig. 4B) in insulin-resistant participants individuals. Adipokine marker Leptin was decreased -7.4% (Fig. 4C) in insulin resistant individuals than in non-insulin resistant individuals. On the other hand, only a 0.6% decrease in HbA1c change in insulin resistant individuals compared to non-insulin resistant controls (Fig. 4E). These findings underscore the efficacy of home-based BW-HIIT in improving cardiometabolic health, particularly in individuals with insulin resistance.

**Fig. 4.**
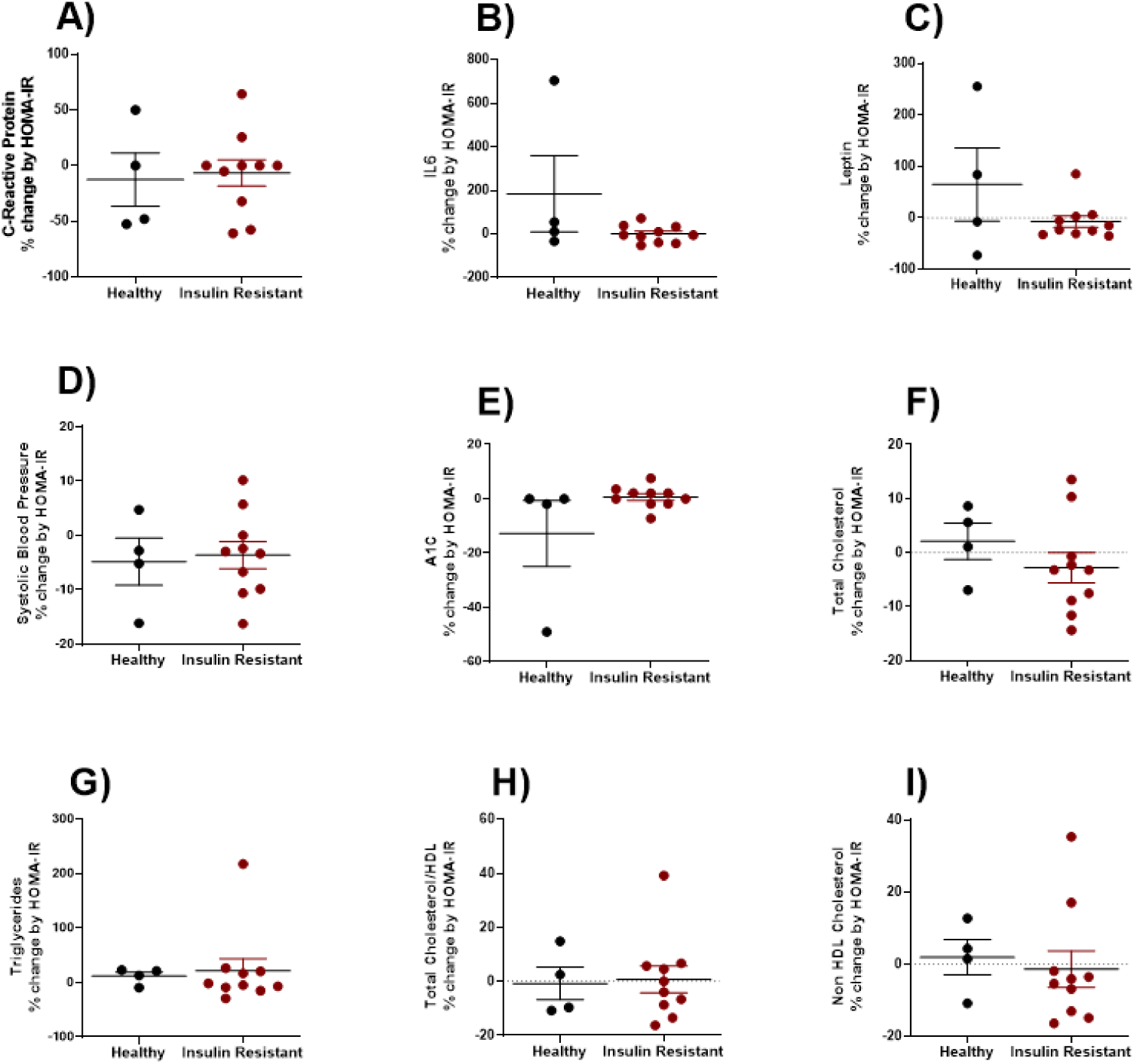
BW-HIIT enhances post exercise cardiometabolic benefits predominantly in insulin resistant individuals with high HOMA-IR. Standard of care and plasma cardiometabolic assessment evidenced the percentage of change sorted by HOMA-IR. Non-insulin resistant (<2 HOMA-IR) vs insulin resistant (>2 HOMA-IR) individuals are presented for the following parameters: A) C-Reactive Protein, B) IL6, C) Leptin, D) Systolic blood pressure, E) A1C (HbA1c), F) Total cholesterol, G) Triglycerides, H) Total cholesterol/HDL, I) Non-HDL cholesterol. All data represented mean+/-SEM. All cytokine levels in insulin resistant (n=10) compared to non-insulin resistant (n=4) were used for independent student t-test analysis.

### 3.4 Plasma lipidomic reveals a beneficial decrease in circulating lipid mediators post exercise in insulin resistant individuals

Lipidomic analysis with exercise in both non-insulin resistant and insulin-resistant individuals illustrate the differential lipidomic profiles between each group (Fig. 5). The Heat Map Analysis, a powerful tool in our arsenal, revealed significant clustering of lipidomic changes between pre- and post-exercise samples in both groups. Insulin resistant individuals exhibited a distinct decreased lipidomic profile compared to their non-insulin resistant counterparts, indicating a more pronounced response to the exercise regimen (Fig. 5A). The EBAM analysis, another high-precision method, identified several lipid species with a significant decrease from pre- to post-exercise almost uniquely in insulin resistant individuals. (Fig. 5B). The PCA loading plot, a robust statistical technique, demonstrated a clear separation between pre- and post-exercise lipid profiles. The post-exercise lipidomic profiles of insulin resistant individuals clustered distinctly from their pre-exercise profiles and the profiles of non-insulin resistant individuals, suggesting specific lipidomic shifts attributable to a decrease in HOMA-IR with insulin resistance and exercise. (Fig. 5C). The SOM plot, a sophisticated visualization tool, highlighted greater variability in the lipidomic data of insulin resistant individuals compared to non-insulin resistant. This variability underscores the heterogeneity in metabolic responses to BW-HIIT exercise within the insulin resistant group (Fig. 5D). Both SPLS and OPLS plots, advanced modeling techniques, confirmed the lipidomic differences between pre- and post-exercise states. These analyses provided robust models for predicting lipidomic changes based on the exercise intervention, particularly in insulin resistant individuals (Fig. 5E, F). The FC plot showed significant upregulation and downregulation of specific lipids in insulin resistant individual’s post-exercise. (Fig. 5G). The linear correlation heat map, a powerful data visualization tool, revealed strong correlations between certain harmful circulating lipid metabolite species decreased in insulin resistant individuals post-exercise, which were less pronounced in non-insulin resistant individuals. This suggests that the effects of modulation of insulin resistance may allow for enhancement of the lipidomic response to exercise (Fig. 5H).

**Fig. 5.**
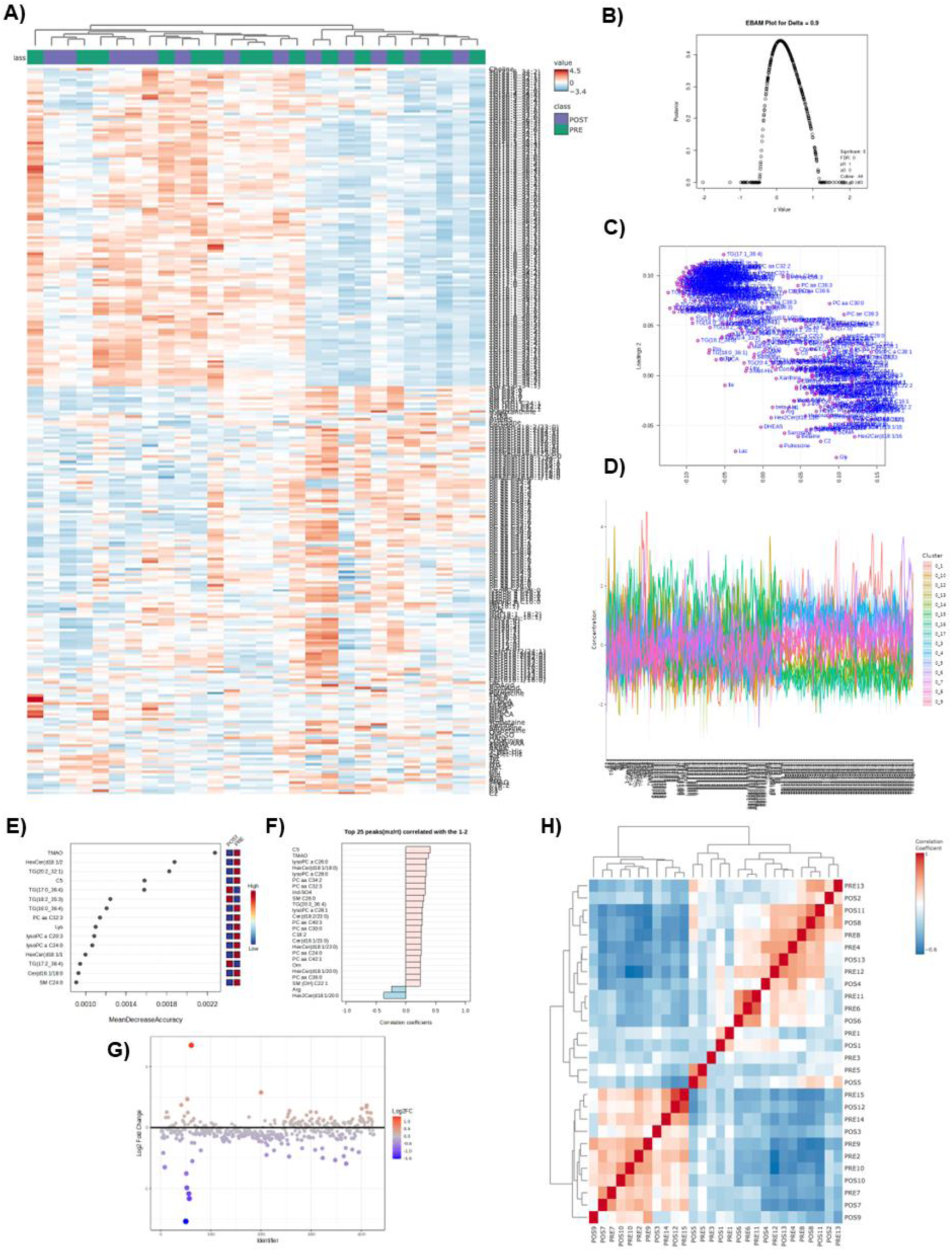
Plasma lipidomics identified overriding clustered differences post-exercise in insulin resistant individuals. A) Heat map of principal changes between pre- vs. post-exercise changes between non-insulin resistant and insulin resistant individuals, B) EBAM plot for delta data presentation, C) PCA loading plot, D) SOM plot for variability in sample data, E) SPLS plot, F) OPLS plot, G) FC plot from post vs. pre-exercise score plot, H) linear correlation heat map between non-insulin resistant vs. insulin resistant individuals after exercise. All data represented mean+/- SEM. N of 4 for non-insulin resistant and 10 for insulin resistant patient groups.

### 3.5 HMGB1 and HOMA-IR Levels follow a similar reduction pattern post-exercise in insulin resistant individuals that is mimicked in mice

Our research has uncovered promising results, graphically illustrating the impact of BW-HIIT of exercise in HOMA-IR and HMGB1 levels. These results, which show a significant metabolic improvement in insulin resistant individuals, are a beacon of promising results. The analyses, including paired and segmented comparisons, delta changes, and comparisons with non-T2D and sedentary chronic T2D individuals (Fig.6), further bolster our optimism. A side-by-side comparison, clustered and segmented, confirms that insulin resistant individuals showed a wider distribution of HOMA-IR than non-insulin resistant control subjects (Fig.6A). Paired comparison of HOMA-IR levels pre- and post-exercise shows a significant decrease in insulin resistant individuals, while non-insulin resistant individuals exhibit less change. This indicates that the improvement of insulin sensitivity is more pronounced in the insulin resistant group post-exercise (Fig.6B). Delta change analysis shows decreased HMBG1 levels post-exercise compared to pre-exercise individuals involved in the BW-HIIT exercise program (Fig.6C). Sub-group analysis shows HMBG1 delta changes are augmented post-exercise (Fig.6D). A sub-analysis identified increased serum HMBG1 levels in sedentary chronic T2D individuals compared to baseline and 6 weeks after exercise of non-T2D (Fig.6E). Mice comparison correlates human findings shown in supplementary Fig. 3. HMGB1 levels show the same results as humans, having a significant decrease in the post-exercise group. These results suggest that BW-HIIT exercise significantly reduces HOMA-IR and HMGB1 levels in insulin resistant individuals, reflecting possible improvement in insulin sensitivity and reduced inflammation. The consistent findings across human and mouse models highlight the potential of BW-HIIT exercise as an effective intervention for metabolic health improvement in insulin resistance with a strong concept of T2D, paving the way for a brighter future in metabolic health research.

**Fig. 6.**
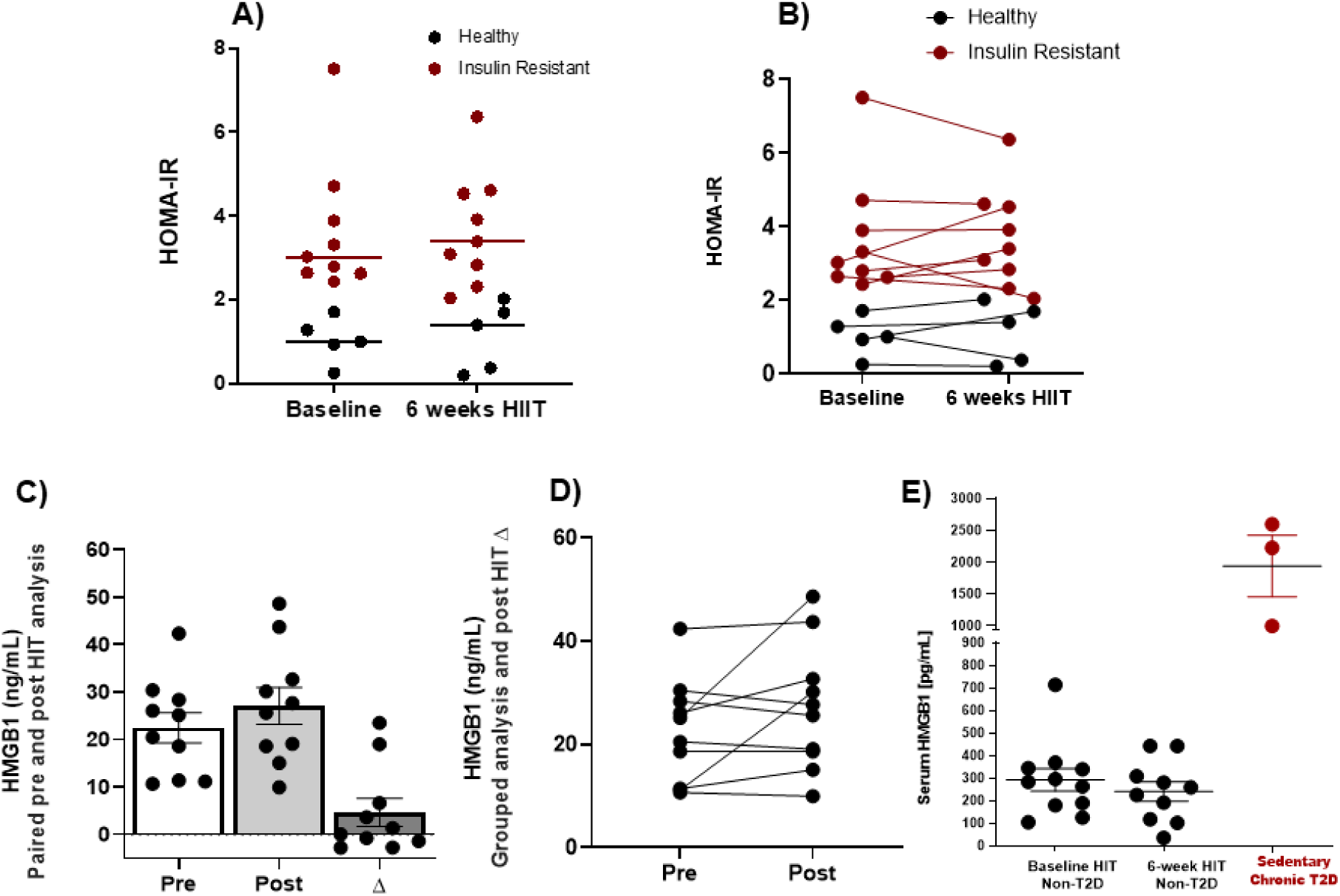
HMGB1 and HOMA-IR levels share associated and potentially causal decreases post-exercise in insulin resistant individuals. A) HOMAIR clustered-segmented side by side comparison (n=5, non-insulin resistant; n=9, insulin resistant), B) HOMA-IR paired comparison(n=5, non-insulin resistant; n=9, insulin resistant), C) Delta change in HMBG1 levels in pre and post-exercise(n of 10 for pre and post exercise), D) HMBG1 levels in pre and post-exercise individually paired data presented (n of 10 for pre and post exercise), E) Serum HMBG1 levels at baseline compared to 6 weeks after BW-HIIT of non-T2D compared to sedentary chronic T2D (n of 11 baseline BW-HIIT; n of 10 for 16 weeks BW-HIIT and n of 3 for Sedentary chronic T2D). All data represented mean+/-SEM. Statistical analysis was performed independently using two-tailed student t-test analysis. Significance levels: *:<0.05, **: 0.001-0.01, ***: 0.0001, ****: <0.0001.

## 4. Discussion

Our study evidenced for the first time (to our knowledge) the effect of decreased circulating HMGB1 in insulin resistant individuals performing maximal effort BW-HIIT as a modality of HIIT due to its cardiometabolic benefits (Fig.1). In addition, we compared these results with animal models following continuous treadmill running to seek for correlation between murine and human results and. We found vast heterogeneity in both models used. Thus, highlighting the translational implications of our findings. However, plasma lipidomics evidenced the major advantages of BW-HIIT in reducing harmful circulating lipid metabolites. Our group conducted this study using the modified HIIT using BW-HIIT in our individuals to highlight the potential cardiometabolic benefits, especially for individuals unable to perform ordinary exercise training regimens using specialized exercise equipment. This underscores the importance of HMGB1 individuals with obesity and T2D individuals since this this population is often sedentary with limited physical capacity. The 6-week BW-HIIT was developed to assess its potential in stimulating changes in metabolic hormones, cardiometabolic markers, cytokine, and lipidomics in insulin resistant adults with obesity. We also developed an 8-week continuous exercise treadmill model for mice, with a focus on to compare findings, focusing on circulating HMGB1. Our findings show: i) anti-inflammatory cytokines IL-10, IL-12p40, and IL-12p70 significantly increased, and the pro-inflammatory cytokines Eotaxin, Il-2, and MIP-2 or CXCL2 significantly reduced in the post-exercise mouse model; ii) Metabolic hormones like Amylin, Glucagon, and Insulin increased in post-exercise human and mouse models; iii) Post-exercise insulin resistant individuals decrease cardiometabolic risk factors including systolic blood pressure, cholesterol, triglycerides, HDL, and Chol/HDL ratio; iv) exercise reduced circulating HMBG1 levels in insulin resistant individuals and exercised mice.

Recent studies suggest that aerobic exercise increases pro-oxidant status more than anaerobic exercise, and aerobic and anaerobic exercise activities improve antioxidants [27, 28]. At the same time, Parker et al. demonstrated that increasing exercise intensity resulted in greater endogenous antioxidant defenses [29–32]. HIIT, a form of exercise that is primarily anaerobic, is characterized by short bursts of intense activity followed by rest periods [33], consequently relying heavily anaerobic energy pathways [34]. This style of training is a well-known stimulus enhancing anaerobic capacity and aerobic fitness [34, 35]. Our research findings, backed by robust evidence, demonstrate that cardiometabolic risk factors, a cluster of factors that increase the likelihood of developing cardiovascular disease and metabolic disorders such as T2D, can be effectively reduced through regular exercise [36, 37]. HIIT’s benefits improve endothelial function, reduce inflammation, and enhance vascular health, thereby lowering blood pressure, improving lipid profiles, and reducing excess body weight [38–41]. Our study on insulin resistance individuals who underwent BW-HIIT demonstrated reduced cardiometabolic risk factors such as systolic blood pressure and cholesterol. Similarly, recent studies have shown positive changes in cardiometabolic risk factors, including systolic blood pressure, in obese individuals undergoing HIIT or a mobile health diabetes prevention trial [42, 43]. In contrast, Bouchonville et al. and their colleagues reported that weight-loss-based lifestyle interventions, not exercise alone, effectively reduced multiple cardiometabolic risk factors in obese older adults [44]. Our study implies that BW-HIIT offers a credible advantage and alternative to traditional styles of exercise training in improving cardiometabolic health and reducing hypertension in adults with obesity.

Our research is directly relevant to the field of exercise physiology and metabolic health. Amylin and Insulin, co-secreted within pancreatic beta cells, are crucial in maintaining blood glucose levels [9, 34, 45]. We observed increased amylin and insulin levels in humans and mice post-exercise, suggesting that BW-HIIT post-exercise directly enhances insulin sensitivity. Conversely, glucagon, a key player in hepatic glycogenolysis and gluconeogenesis during exercise, showed elevated plasma levels in post-exercise humans and mice. Ghrelin, a regulator of appetite, energy balance, glucose metabolism, and insulin sensitivity, was also affected. Our study revealed that post-exercise improves metabolic hormones and increases ghrelin levels in insulin-resistant individuals. These findings parallel those of a recent study that showed the effect of lifestyle intervention in individuals with obesity and insulin resistance, which showed a 23.6 % reduction in Ghrelin levels after four months of intervention [46].

Our study demonstrated that HMBG1 levels can be reduced post-exercise in insulin resistant adults. This finding is significant and holds immense potential as it provides a potential mechanism for the beneficial effects of BW-HIIT exercise on insulin resistance. HMBG1, a key DAMP-released protein, triggers inflammation that can infiltrate into tissue, a process strongly associated with insulin resistance [16, 47, 48]. Importantly, our findings align with those of Gmiat et al., who observed decreased HMBG1 levels after 12 weeks of Nordic walking with vitamin supplementation by older women, suggesting that regular exercise may diminish the response in non-insulin resistant adults [49, 50]. However, in contrast, the HMBG1, the receptor for advanced glycation end products and nucleosomes, increases after a marathon [51, 52]. These results underscore the high potential but distinct acute vs. chronic effect of exercise training that can modulate circulating HMGB1, thereby serving as a valuable biomarker in both non-insulin resistant and clinical conditions. This potential opens exciting avenues for further research and clinical applications, especially for our group [53]. These findings are also contrasted with the effect of acute vs chronic responses to HMGB1[54]. These findings, which echo those of a recent study on HMBG1 levels pre and post marathon, suggest potential beneficial effects of marathon running on the body. The HMGB1 level was analyzed 10-12 weeks prior, 1-2 weeks before, immediately, 24 h, 72 h, and 12 weeks post-race. The results demonstrated a significant increase in HMGB1 from pre- to immediate post-race, followed by a return to baseline within 24-72 h. It’s worth noting that strenuous exercise induces distinct changes in circulating HMGB1, and marathon running can trigger extensive responses from the body, potentially leading to beneficial effects, as observed in these subjects [51, 52]. These findings direct us to our future studies to investigate recruiting and retaining more chronic T2D individuals and provide them with tailor-made supervised BW-HIIT exercise training that could improve their glycemic control, thereby retaining nuclear HMGB1 signals. These implications for future research highlight the need for further investigation into the acute and chronic effects of exercise on HMGB1 levels and the potential benefits of exercise on insulin resistance.

We are mindful of the limitations of our study. First, we must include that all individuals were instructed remotely to perform BW-HIIT workouts three days a week over six weeks. This remote instruction could have led to variations in the intensity and adherence to the training program [55]. However, due to the limitation of cohort individuals, this study reveals the main effect of systemic HMBG1 in non-insulin resistant vs insulin resistance on exercise levels. Analyzing HMBG1 changes on pre- and post-exercise training is also challenging since some of our individual’s showed variability in the extent of the exercise they performed [56]. This study aimed to evaluate HMGB1 as a useful biomarker, which will be ideal for assessing anti-hyperglycemic treatment efficacy in future studies [57]. Additionally, the study’s smaller size limits the generalizability of the findings. For this study, we focused strongly on retaining a group of individuals with insulin resistance. Due to BW-HIIT being performed remotely, research members tried to check-in with individuals if their WAT indicated they were not performing the workouts. In the future, prospective studies will aim to recruit a higher number of subjects that are sex and age-matched within non-insulin resistant and T2D individuals with the presence and absence of diabetes-associated complications and provide them with tailor-made supervised BW-HIIT exercises. Furthermore, the study did not include a control sedentary group, which could have provided a better comparison of the effects of HMGB1 and BW-HIIT exercise. However, a pilot study by Goh et al. in 2020 examined the impact of concurrent high-intensity aerobic and resistance exercise on releasing alarmins such as HMBG1 and inflammatory biomarkers in non-insulin resistant young men and found elevated plasma HMBG1 levels in immediately after (Post), 30 min (30 min) after exercise each week, and 24 h after the final exercise session in week 3 (24 h). Importantly, we cannot definitively exclude sex differences in the effects of exercise training on the HMBG1 level. These limitations should be considered when interpreting the study’s results.

In the future, we plan to extend the study to include age matched chronic T2D patients beyond six weeks, using supervised training. We will measure HMGB1 levels in blood and blood-derived extracellular vesicles to understand how HMGB1 is released in insulin-resistant patients and those with diagnosed T2D. This next step is not only crucial but also holds promise as it will help us understand the potential of HMBG1 in regulating insulin secretion in response to exercise training. These future mechanistic studies will be pivotal in determining the role of HMBG1 in our understanding of diabetes management and obesity treatment. Furthermore, additional research is needed to develop culturally adaptable versions of vigorous-intensity interval training using one’s own bodyweight. Such studies will promote inclusive training protocols for diverse genders, demographics, and minorities in New Mexico and the US population and improve their quality of life.

## 5. Conclusions

The current study demonstrated that 6-weeks of bodyweight high intensity interval training improves cardiometabolic health, anti-inflammatory markers, metabolic hormones, and insulin sensitivity in human and preclinical exercise mouse models. Correlatively, changes in circulating HMBG1 using BW-HIIT exercise training make HMGB1 a suitable marker for metabolic disease potentiating its role beyond an alarmin. Further studies are needed to confirm these effects and to elucidate the underlying physiological mechanisms. Overall, these experiments reinforce the potential of BW-HIIT exercise interventions in insulin resistant adults to prevent development of T2D.

## Supplementary Material

### 2.5 Supplemental methods

All animal experiments were approved by the ethics committee of the University of New Mexico (UNM) Institutional Animal Care and Use Committee (IACUC) and Animal Welfare Committee. All live animal studies were conducted ethically, following relevant guidelines and regulations at the University of New Mexico under protocol number 23-201405-HSC. All animals were housed under the Animal Resource Facilities (ARF) in UNM. At the end of the experimental protocol, the animals were euthanized using a dose of 0.01 mL/g of Ketamine/Xylazine. Harvested tissues included blood, liver, and muscle. In supplementary figure 3b, mice were considered as insulin resistant mice due to the implementation of a High-fat diet (HFD) (D12492, Research Diets) that was administrated starting at 6 weeks of age before exercise training. For T2D model development, an HFD (D12492, Research Diets) was administrated at 6 weeks of age. At 8 weeks of age, an intraperitoneal (IP) injection of Streptozotocin (STZ) was given for 5 consecutive days at a dosage of 25 mg/kg to the mice because this induces Hyperglycemia and insulin Resistance. Following the final STZ injection, the mouse model was monitored and allowed to evolve over 10 weeks. These mice were placed in treadmills to run daily for 1hr at a velocity of 80 meters/min for 4 weeks. In supplementary figure 3c, a non-exercise and HFD (D12492, Research Diets) was administrated at 6 weeks of age to HMGB1 Flox mouse models that were developed as previously described. A tamoxifen (TMX) injection was administered to the mice at 6 weeks of age by intraperitoneal (IP) injection at a dose of 1 mg/kg over 10 days to lack the inducible mechanism for HMGB1 knockdown. For the second group, the same procedure was followed; after a 7-day post-TMX injection period, mice were administered a low dose of Streptozotocin (STZ) via IP for 5 consecutive days at a dosage of 25 mg/kg to make them T2D. Their state was monitored and allowed to evolve over 10 weeks. For supplementary figure 3, an ELISA kit (MyBioSource, MBS701378) was used to measure the levels of HMGB1 in our samples.

**Supplementary Fig. 1.**
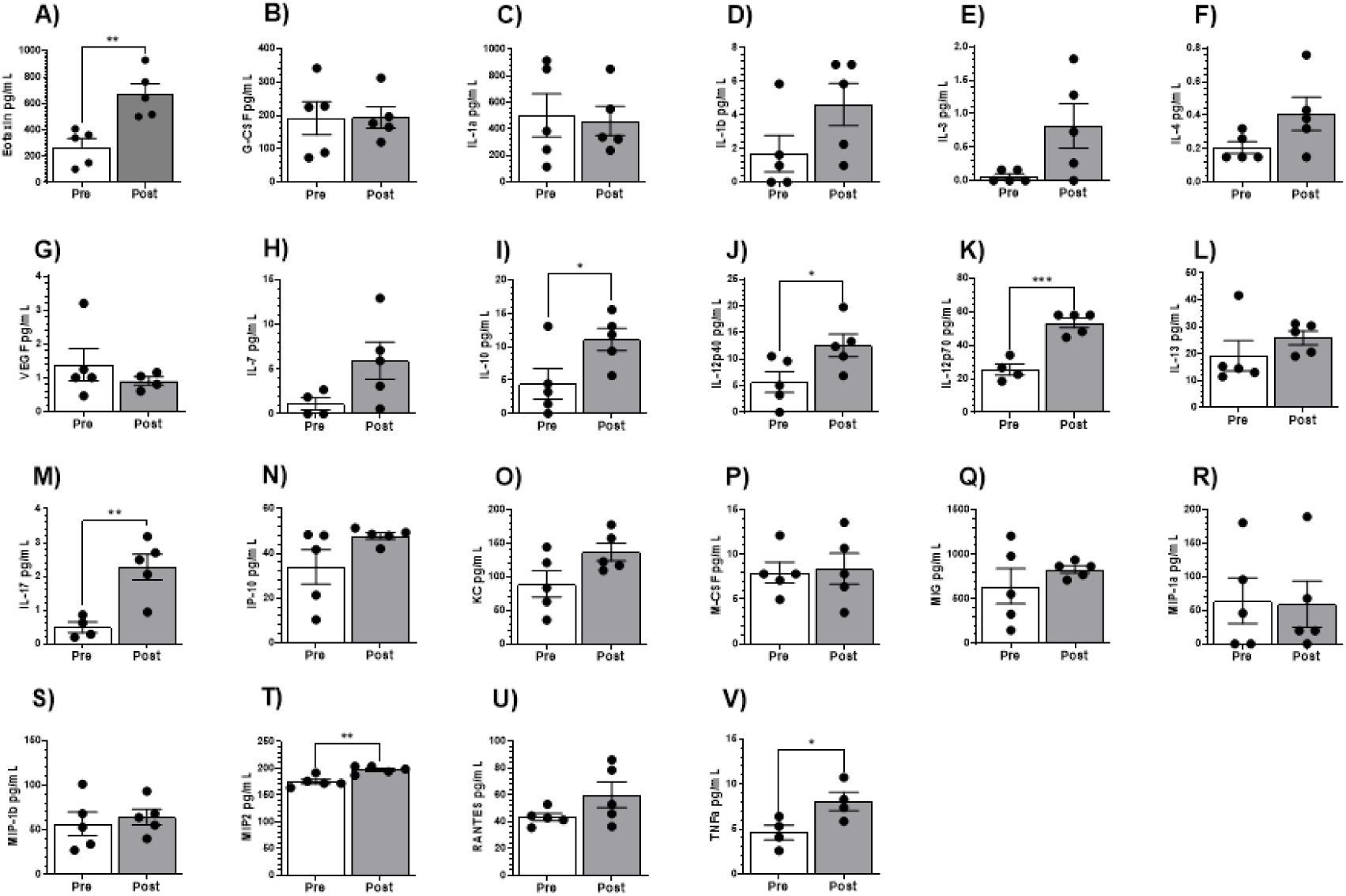
Unmatched plasma levels of cytokine panel in pre and post exercise in mice. A) Eotaxin, B) G-CSF, C) IL-1a, D) IL-1b, E) IL-3, F) IL-4, G) VEGF, H) IL-7, I) IL-10, J) IL-12p40, K) IL-12p70, J) IL-13, M) IL-17, N) IP-10, P) M-CSF, Q) MIG, R) MIP-1a, S) MIP-1b, T) MIP2, U) RANTES, and V) TNFa. All data represented mean+/-SEM. All cytokine levels in post-exercise(n=5) compared to pre-exercise (n=5) were used for independent student t-test analysis. Significance levels: *:<0.05, **: 0.001-0.01, ***: 0.0001, ****: <0.0001.

**Supplementary Figure 2.**
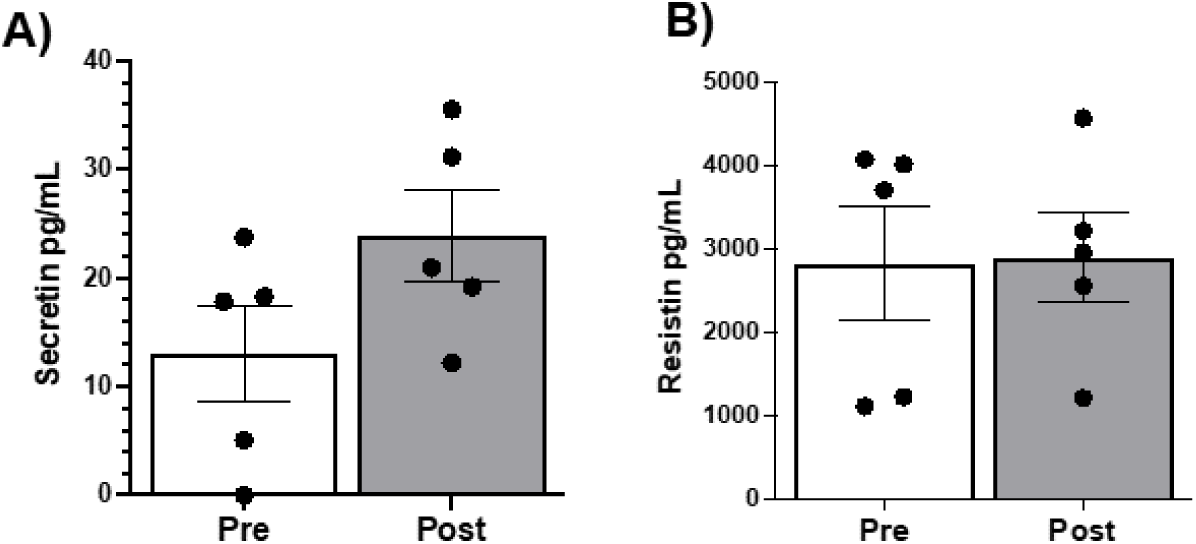
Unmatched plasma levels of metabolic hormones in pre and post exercise samples from mice. Analysis shows comparison between pre and post treadmill at levels of Secretin (A) and Resistin (B). All data represented mean+/-SEM. All cytokine levels in post-exercise(n=5) compared to pre-exercise (n=5) were used for independent student t-test analysis. Significance levels: *:<0.05, **: 0.001-0.01, ***: 0.0001, ****: <0.0001.

**Supplementary Figure 3.**
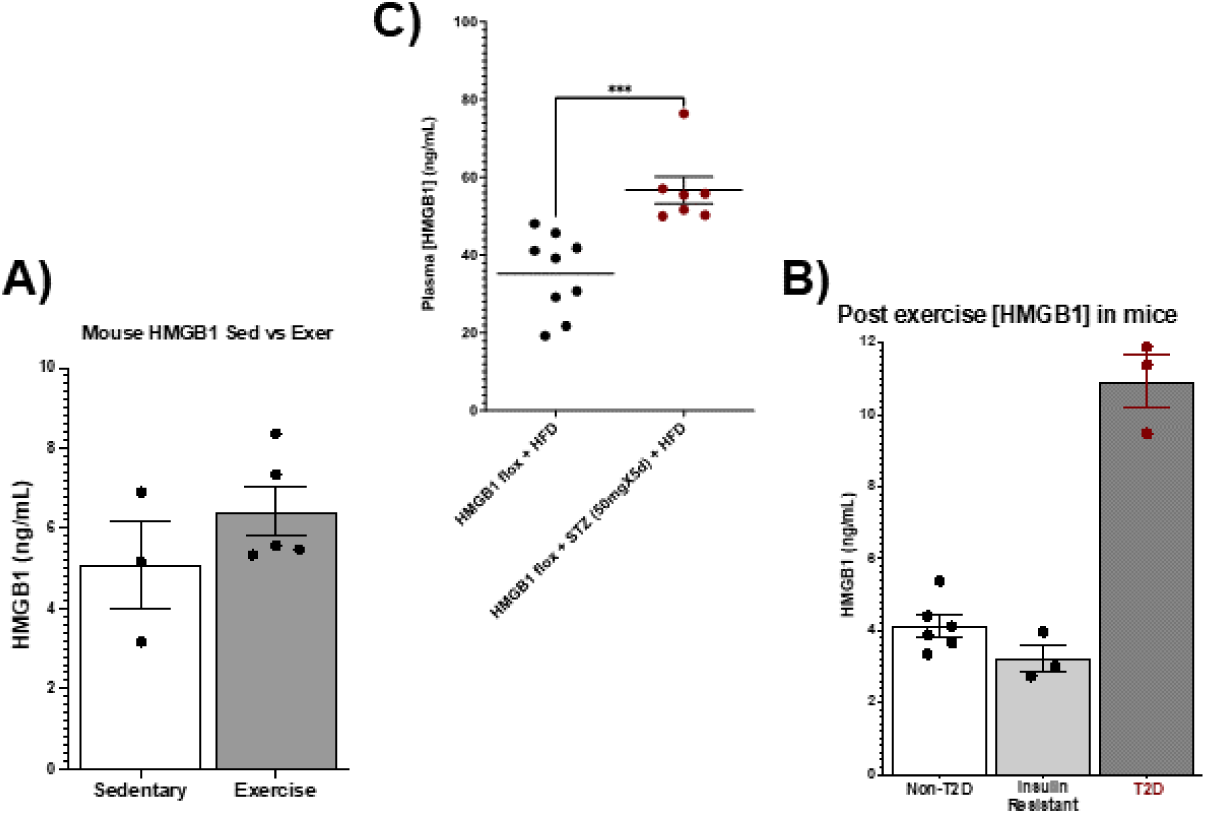
Comparison of HMBG1 levels in distinct mouse models of Insulin Resistance and T2D. A) HMBG1 levels in sedentary (n=3) vs. exercise mice(n=5); B) HMBG1 levels in post-exercise non-T2D (n=6), insulin resistance(n=3), and T2D mice (n=3); and C) HMBG1 levels in HMBG1 Flox mice fed with HFD and HMBG1 Flox mice fed with HFD and STZ. All data represented mean+/-SEM. Independent student t-test analysis performed between groups. Significance levels: *:<0.05, **: 0.001-0.01, ***: 0.0001, ****: <0.0001.

## Availability of Data and Materials

The data that support the findings of this is available in the MENDELEY DATA repository: Mota Alvidrez, Roberto (2024), “High-Intensity Interval Training Decreases Circulating HMGB1 in Insulin Resistant Individuals; Plasma Lipidomics Identifies Associated Cardiometabolic Benefits”, Mendeley Data

## Author Contributions

GMB, PP: Writing – review & editing, Writing – original draft, Software, Methodology, Investigation, Formal analysis, Data curation. QJ, GFB, HZ: Validation, Resources, Methodology, Formal analysis, Data curation. FA: Visualization, Validation, Supervision, Resources, Methodology, Funding acquisition, Conceptualization. RIMA: Writing – review & editing, Writing – original draft, Visualization, Validation, Supervision, Software, Resources, Project administration, Methodology, Investigation, Funding acquisition, Formal analysis, Data curation, Conceptualization. All authors contributed to editorial changes in the manuscript. All authors read and approved the final manuscript.

## Ethics Approval and Consent to Participate

The Ethics Committee of The University of New Mexico (UNM) Institutional Review Board (IRB) Main Campus approved the research protocol with reference number 21-319. All individuals provided signed informed consent. All animal experiments were approved by the ethics committee of the University of New Mexico (UNM) Institutional Animal Care and Use Committee (IACUC) and Animal Welfare Committee. All live animal studies were carried out ethically, following relevant guidelines and regulations at the University of New Mexico under protocol number 23-201405-HSC.

## Acknowledgment

We gratefully acknowledge the assistance and instruction from Dr. Fabiano Amorim, his team for the help with the human samples used for this study. We also acknowledge the assistance of Eve Technologies Corporation in processing the human and mouse samples. We would also like to thank Dr. Changjian Feng and his lab members for their help with the lipidomic studies and analysis.

